# biHash: bisulfite-aware recursive nucleotide hashing

**DOI:** 10.64898/2026.10.19.760264

**Authors:** Johnathan Wong, Inanc Birol

## Abstract

Bisulfite sequencing (BS-seq) measures cytosine methylation on the DNA at single-base resolution, but converts unmethylated cytosines to thymines, disrupting conventional strand complementarity and reducing sequence complexity. These pose a fundamental challenge to efficient sequence comparison and indexing tasks. Although *k*-mer hashing is foundational to bioinformatics for enabling such high-throughput operations, generic algorithms for string hashing are poorly equipped to handle the reduced-complexity alphabet and sequence ambiguity created by this conversion. We therefore require fast *k*- mer hashing that directly recognizes bisulfite-compatible sequence states. Here, we present biHash, a rolling hash algorithm tailored for bisulfite-sequenced data. Using context-specific dinucleotide encodings, biHash maps designated bisulfite-compatible sequence variants to the same hash value while retaining context-specific specificity. When hashing adjacent *k*-mers, biHash performs ∼13 times faster than generic string hashing algorithms.

## Introduction

Analyzing the epigenetic landscape provides biological insight that the DNA sequence alone cannot capture. DNA methylation, particularly at the fifth carbon position of cytosine (5-methylcytosine, 5mC), is a fundamental epigenetic modification that is involved in transcriptional silencing, genomic imprinting, X-chromosome inactivation, and the maintenance of genome stability (Moore, Le, and Fan 2013). Together with other modifications like 5-hydroxymethylcytosine (5hmC), these marks form intricate, genome- wide patterns that are associated with cell specialization and developmental trajectories (Bai *et al*. 2025, Honig and Murrell 2025). Clinically, altered epigenetic patterns can provide signatures of disease; for example, hypermethylation of tumor suppressor gene promoters is a well-established hallmark of early-stage cancer (Esteller 2006).

Bisulfite sequencing (BS-seq) is widely used to profile cytosine modifications at single- base resolution (Chen *et al*. 2025), although standard BS-seq cannot distinguish 5- methylcytosine from 5-hydroxymethylcytosine (Huang *et al*. 2010). The process involves treating DNA with sodium bisulfite, which chemically deaminates unmethylated cytosines into uracils (read as thymines during sequencing) while leaving methylated cytosines intact. In fully converted regions, this reduces the sequence alphabet from four bases to three ∑ = {A, G, T}. Because conversion affects strands asymmetrically, BS-seq reads break the canonical Watson-Crick complementarity (Tran *et al*. 2014). This reduced complexity and strand asymmetry complicate their alignment to the reference genome.

BS-seq aligners such as Bismark and BS-Bolt address strand asymmetry using *in silico* conversion (Krueger and Andrews 2011, Farrell *et al*. 2021). They represent the reference strands using C→T and G→A conversions, then compare correspondingly converted reads against these representations. How these comparisons are performed depends on the tool and library type. Bismark, for example, considers two strand alignments for directional libraries and four for non-directional libraries. Abismal instead uses a two-letter encoding to identify candidate mapping positions, which are subsequently evaluated against the genomic sequence to resolve ambiguous matches (de Sena Brandine and Smith 2021).

Efficient sequence comparison depends in part on how short sequence fragments, or *k-* mers, are represented and queried. Hashing provides one way to index and compare these fragments. Rolling hashes are particularly efficient for consecutive *k*-mers because each value can be updated from the preceding window (Mohamadi *et al*. 2016). Degenerate encodings can additionally assign selected sequence variations to the same hash value, allowing them to be considered together during *k*-mer matching (Wong *et al*. 2026).Despite advances in *k*-mer hashing, we are not aware of a rolling hash algorithm designed to encode bisulfite-compatible sequence states directly. Existing nucleotide hashing algorithms generally operate on regular DNA sequences and do not, without additional conversion steps, assign bisulfite-compatible variants to the same hash value. Here, we present biHash, a bisulfite-aware rolling hash algorithm that utilizes context-specific dinucleotide encodings to support matching between selected reference and bisulfite-derived *k*-mers.

## Methods

biHash builds on ntHash, a rolling hash algorithm for DNA/RNA sequences, and adapts it to the constraints of bisulfite-sequencing data (Mohamadi *et al*. 2016, Kazemi *et al*. 2022). To address the non-complementary sequences introduced by chemical conversion, biHash employs a dinucleotide hashing scheme designed to assign compatible reference and bisulfite-derived sequences to the same hash values (Fig. 1a–d). Each scheme is tailored to a specific cytosine dinucleotide context—CG, CC, CT, or CA—as consolidating them into a single scheme would introduce excessive degeneracy and compromise hash specificity. We also implemented standalone mononucleotide modules for C→T and G→A conversion, as well as a module that combines both conversions. By leveraging the degeneracy logic established in aaHash, these modules provide alternatives to *in silico* sequence conversion.

**Fig 1.**
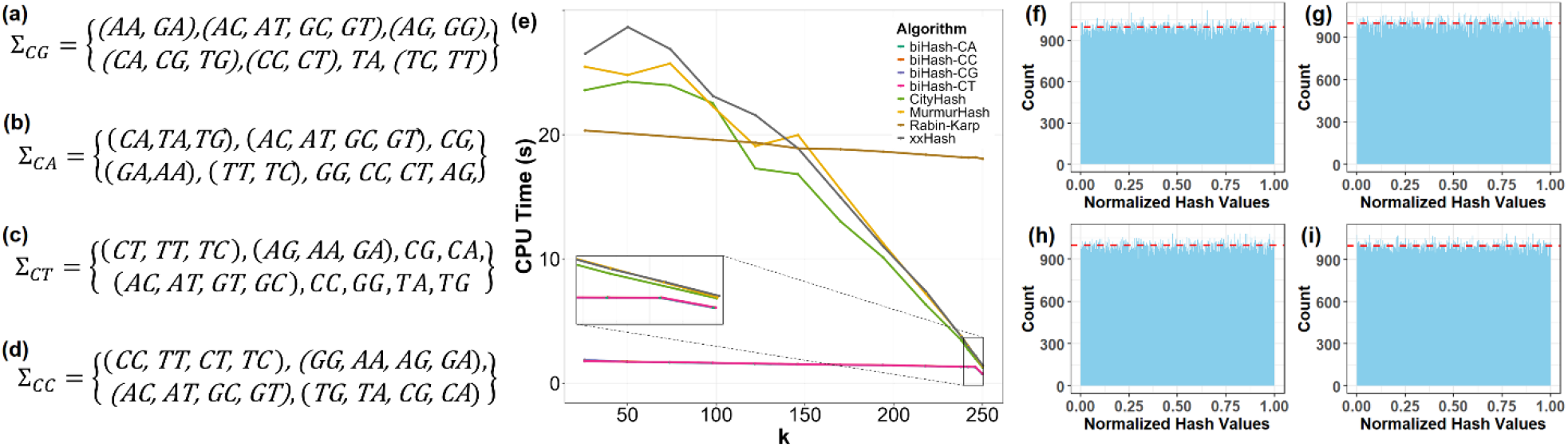
Hashing Schemes and Performance of biHash. (a-d) biHash dinucleotide hashing schemes by CG, CA, CC, and CT methylation context. Within each scheme, dinucleotides enclosed in parentheses are hashed to identical values. This effectively groups bisulfite-converted variants with their original genomic forms, enabling consistent *k*-mer matching regardless of methylation state. (e) Run time for hashing 1,000,000 × 250 nucleotides long sequences with *k*-mer lengths from 26 to 250. biHash outperforms all other hashing methods when computing one or more adjacent *k*-mers, achieving up to ∼13X speedup. (f-i) Histogram of 1,000,000 100-mer hashes generated by the dinucleotide modules of biHash from a random nucleotide sequence of length 1,000,099. The red dashed line indicates the average number of hashes in a bin (1,000). The hash values were normalized by dividing the hash values by 2^64^ − _1_, the largest 64-bit integer, and plotted on the histogram with bin size of 1,000. The mean and standard deviation of the bin counts are 1,000.0±32.01, 1,000.0±31.11, 1,000.0±30.87, and 1,000.0±32.53 for CG, CA, CC, and CT modules, respectively, demonstrating the empirical uniformity of biHash.

Equation (1) defines the initial (base) biHash value *H*(*s*_0_) for a *k-*mer *s*_0_ of even length *k*. The lookup table *H*_*m*_ maps each dinucleotide to a 64-bit integer under context-specific scheme *m*.

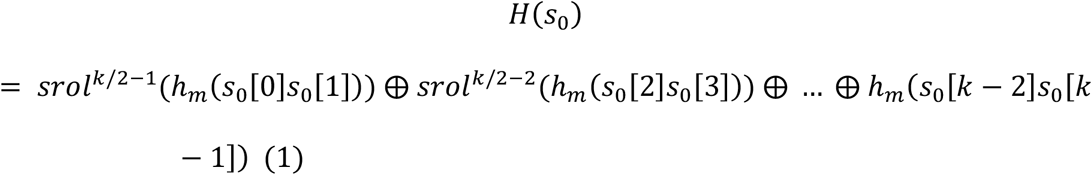

Equation (2) shows the s*rol* operation, which denotes a cyclical bit-rotate-and-swap, is shown in equation (2). It applies *swap*_*bits* function between the 31- and 33-bit segments following a bitwise left-rotate operation, *rol*. This split-rotation operation increases the rotation period from 64 to 1,023, as in ntHash v2.3.0.

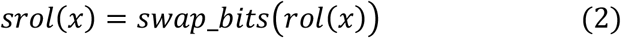

The recursive formula of biHash is shown in Equation (3), where *s*_*i*_ denotes the *i*th *k-*mer in the sequence.

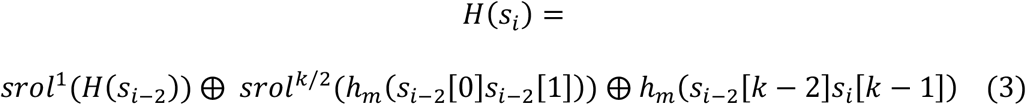

Each dinucleotide-aware rolling update advances the window by two bases, from position i to i+2. To generate a hash for every *k*-mer, biHash maintains separate rolling states for windows beginning at even and an odd positions.

The standalone mononucleotide modules utilize the base and recursive formulas of ntHash v2.3.0, with aaHash-style (v1.1.0) lookup table mappings (Wong *et al*. 2023). The two 3-base modules treat either C and T or G and A as equivalent; the 2-base module treats both pairs as equivalent.

We compared biHash’s runtime and peak memory usage with CityHash, MurmurHash, xxHash, and Rabin–Karp (Supplementary Table S1). We also assessed the distribution of biHash hash values. biHash is implemented within the btllib common code library v1.8.0 (Nikolić *et al*. 2022). All benchmarks were performed using a single thread on a server-class system with 144 Intel(R) Xeon(R) Gold 6254 CPU @ 3.1GHz with 2.9TB RAM. Statistical tests were conducted using Python (v3.10.12) and SciPy (1.11.3) (Virtanen *et al*. 2020).

## Results

To evaluate biHash’s performance in hashing consecutive nucleotide *k*-mers, we hashed 1,000,000 simulated DNA sequences, each 250 nucleotides long, using three biHash module types, four comparators: CityHash, MurmurHash, xxHash, and Rabin-Karp, the last one a cyclical rolling hash algorithm. To simulate bisulfite converted inputs, the sequences evaluated by the comparator algorithms were subjected to *in silico* C→T and G→A conversions. Overall, all three biHash module types consistently outperformed the comparator algorithms. The dinucleotide modules (CG, CA, CC, CT) of biHash were faster than all competitors when hashing at least one *k*-mer, and achieved up to ∼13-fold speedup over CityHash, the next fastest algorithm (1.88s, 1.78s, 1.80s, 1.78s, respectively, vs 23.58s) when hashing 225 consecutive 26-mers (Fig. 1e). Likewise, the 3-base (using both C→T and G→A conversion modules) and 2-base conversion modules performed with similar efficiency, demonstrating ∼12X and ∼22X speedups (2.04s and 1.08s), respectively (Supplementary Figs. S1 and S2).

Next, to assess the quality of biHash values, we measure their uniformity and independence with several methods used in our previous work. A uniform distribution in the hash space helps reduce collisions, an important quality of ideal hash functions. We first evaluated the biHash hash values by plotting a histogram of 1,000,000 normalized hash values derived from 100-mers of a random nucleotide sequence (Fig. 1f-i). Across 1,000 bins, the four hashing schemes within the dinucleotide module yielded means and standard deviations of 1,000.0±32.01, 1,000.0±31.11, 1,000.0±30.87, and 1,000.0±32.53, respectively, closely approximating the expected mean of 1,000 counts per bin. Similarly, the 3-base (C→T and G→A) and 2-base conversion modules exhibited comparable bin count distributions, yielding means and standard deviations of 1000±31.91, 1000±32.52, and 1000.0±31.89 (Supplementary Figures S3-5), respectively.

We also used a Kolmogorov–Smirnov (K-S) test to compare each module’s hash-value distribution with a uniform distribution. The null hypothesis is that the values follow a uniform distribution. The tests did not reject uniformity for any module at *α =* 0.05 (Supplementary Table S2).

We then applied the Box-Muller transformation to pairs of adjacent biHash hash values to examine their distribution and dependence. The transformation converts two uniform, independent inputs, *U*_0_ and *U*_1_, into two independent, normally distributed outputs, *Z*_0_ and *Z*_1_. Adjacent *k-*mer hashes of each module were selected as the input. We tested the normality of the two transformed distributions using the Shapiro-Wilk (SW) test. The test reported *α =* 0.05 SW test statistic of approximately 0.99 and did not reject the normality (Supplementary Table S3). A Pearson correlation coefficient was then computed between the *Z*_0_ and *Z*_1_ distributions. The resulting correlations were all consistent with zero (Supplementary Table S4). These tests found no clear departure from the expected behaviour of transformed independent uniform values, but do not prove that the original hashes are uniform or independent. While cyclical recursive n-gram hashing is not formally uniform in a strictly theoretical sense, these empirical results support the use of biHash in BS-seq applications.

Finally, we compared the peak memory usage of each hashing algorithm. While memory usage remained stable across varying *k*-mer lengths, biHash demonstrated a significantly lower measured peak memory usage, approximately 37.5% less (5.76MB vs. 9.22MB) than traditional hashing functions like CityHash, MurmurHash, xxHash, and Rabin-Karp. This proportional difference reflects the advantage of the degeneracy in hash space since traditional methods require maintaining multiple *in silico* bisulfite-converted copies of each sequence, which increases RAM usage (Supplementary Table S5).

These performance gains reflect the biHash architecture, which was specifically engineered to address the sequence asymmetries of bisulfite-converted data. By leveraging dinucleotide context to partition the alphabet into degenerate groups, it allows specified methylated and unmethylated sequence variants from the same region map to identical hash values. This encoding strategy allows the algorithm to treat the specified bisulfite-converted sequences and their reverse complements as equivalent states, effectively restoring the symmetry lost during chemical treatment (Supplementary Figure S6). In this framework, we envision a *k*-mer design composed of an odd number of dimers, which positions the target dinucleotide at the center. This design utilizes the central dimer to represent the methylation-relevant context, while the flanking dimers provide sequence context for read mapping.

biHash is a specialized nucleotide hashing algorithm for BS-seq that outperforms the tested hashing algorithms when hashing consecutive nucleotide *k*-mers, a common operation in bioinformatics. Furthermore, its encoding natively accommodates the *in silico* C→T and G→A conversions needed to work with BS-seq data. By combining these specialized matching capabilities with an order of magnitude improvement in hashing speed, we anticipate that biHash will streamline the development of BS-seq analysis software.

## Acknowledgements

None.

## Data availability

Information about the data used to benchmark biHash and its comparators can be found in Supplementary Note S1.

