## Supplementary Information for "biHash: bisulfite-aware recursive nucleotide hashing"

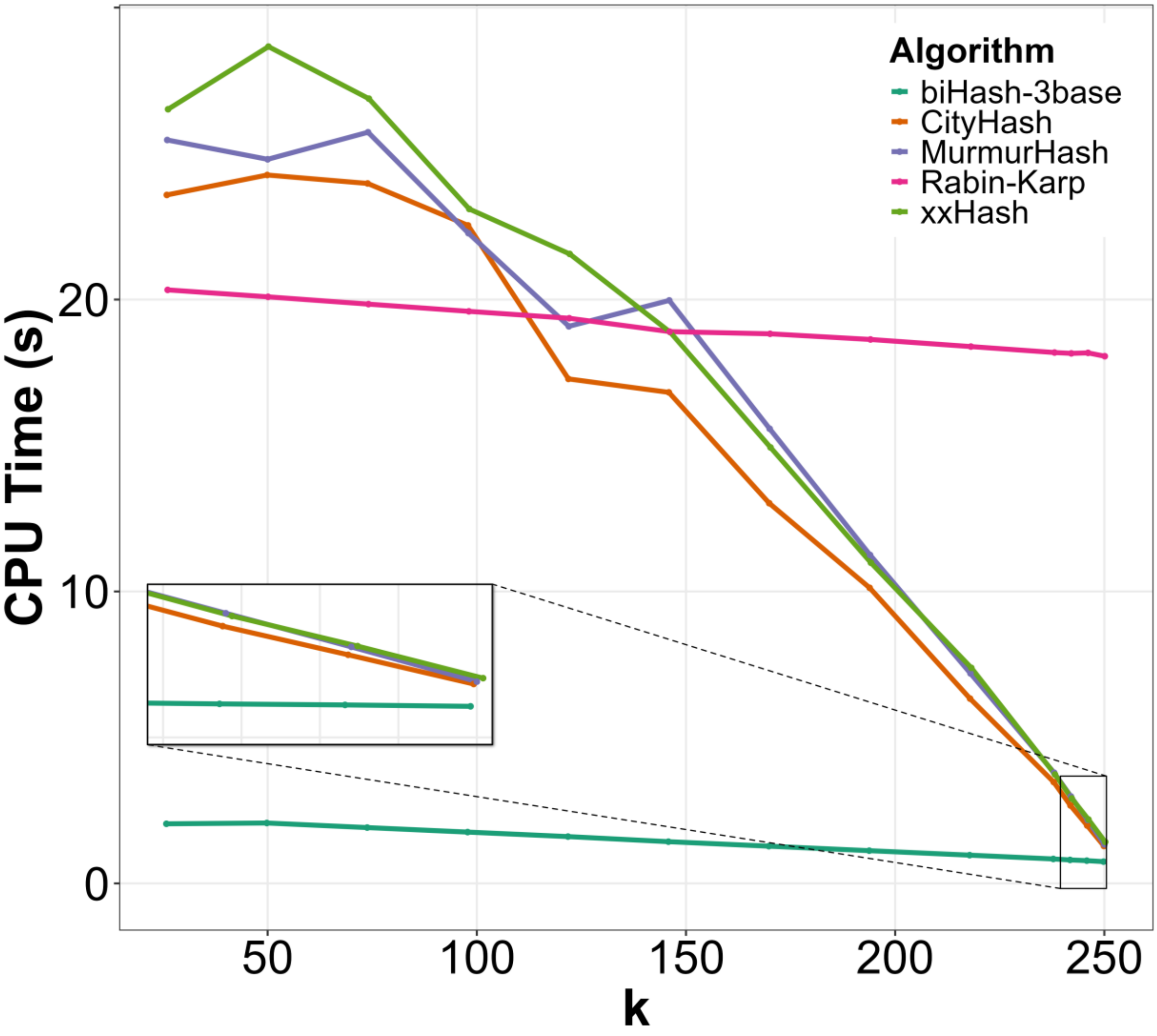

**Supplementary Figure S1. Run time for hashing 1,000,000 x 250 nucleotides long sequences with *k*-mer lengths from 26 to 250 using biHash 3-base module against comparators.** biHash 3-base module, including run time from both C→T and G→A conversion modules, outperforms all other hashing methods when computing more than 1 or more adjacent *k*-mers.

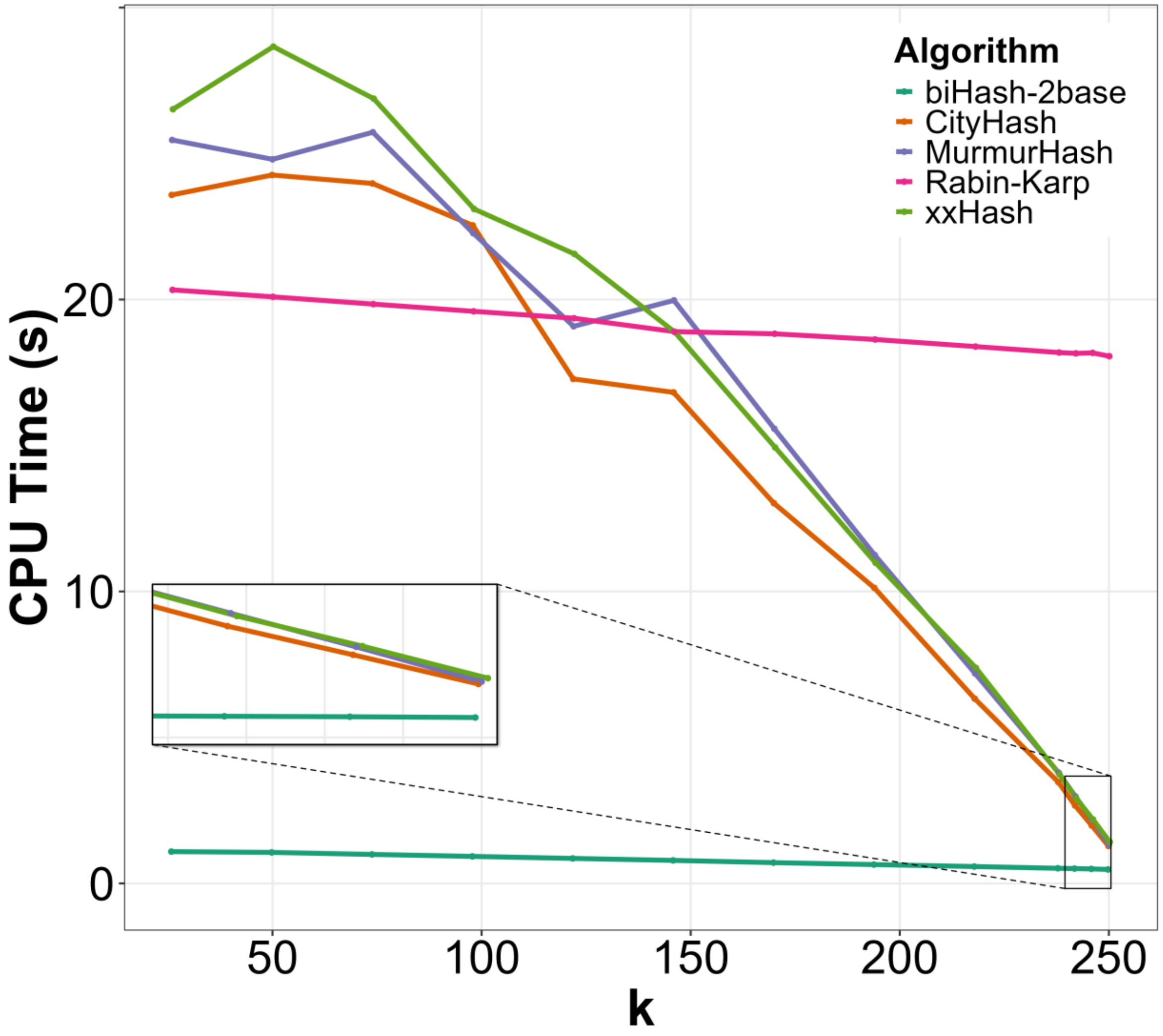

**Supplementary Figure S2. Run time for hashing 1,000,000 x 250 nucleotides long sequences with *k*-mer lengths from 26 to 250 using biHash 2-base module against comparators.** biHash 2-base module outperforms all other hashing methods when computing more than 1 or more adjacent *k*-mers.

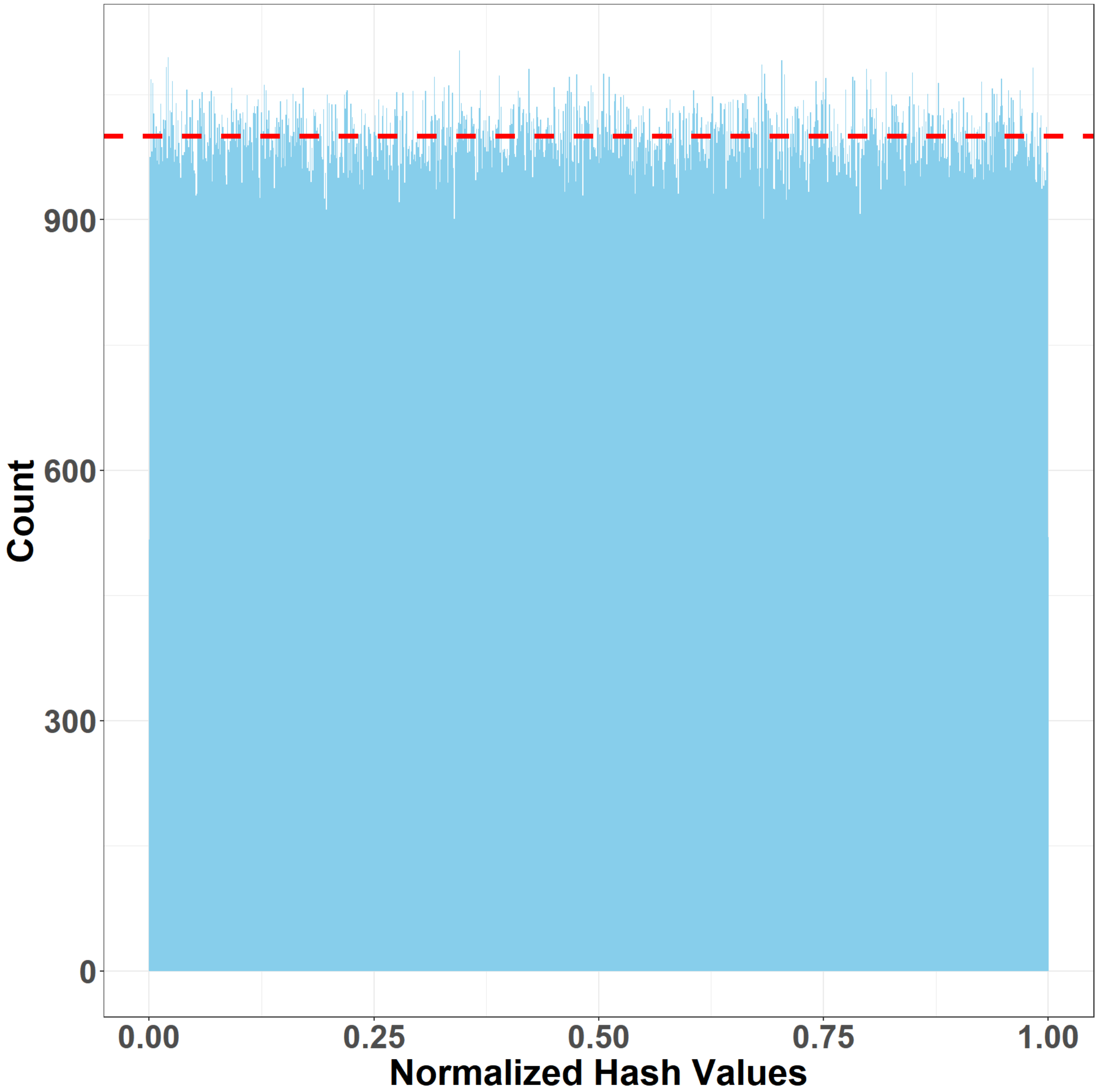

**Supplementary Figure S3. Histogram of 1,000,000 100-mer hashes generated by 3-base C→T conversion module of biHash from a random nucleotide sequence of length 1,000,099.** The red dashed line indicates the average number of hashes in a bin (1,000). The hash values were normalized by dividing them with $2^{64} - 1$, the largest 64-bit integer, and plotted on the histogram with bin size of 1,000. The mean and standard deviation of the bin counts are 1000.0±31.91, demonstrating the uniformity of biHash.

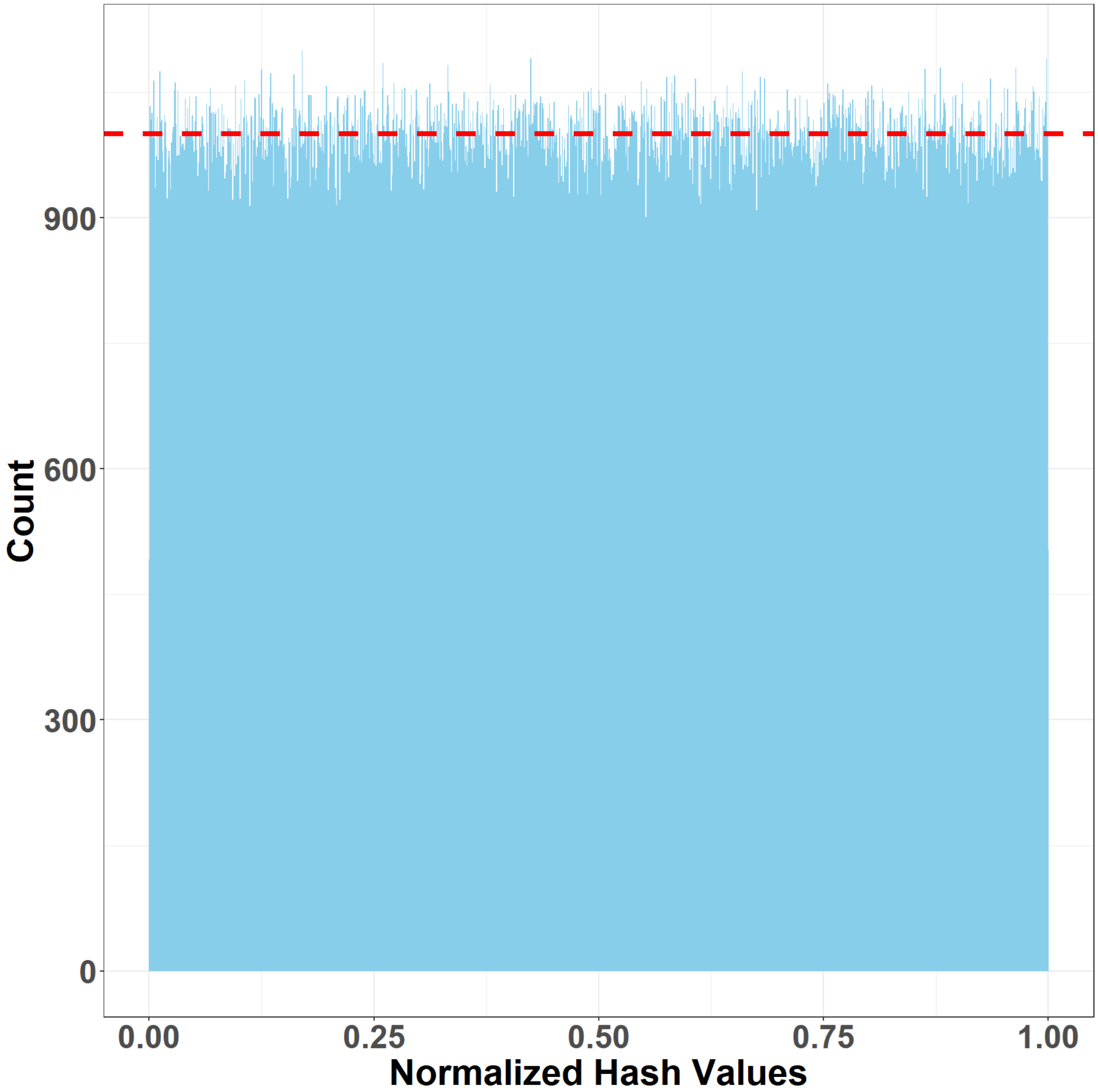

**Supplementary Figure S4. Histogram of 1,000,000 100-mer hashes generated by 3-base G→A conversion module of biHash from a random nucleotide sequence of length 1,000,099.** The red dashed line indicates the average number of hashes in a bin (1,000). The hash values were normalized by dividing them with $2^{64} - 1$, the largest 64-bit integer, and plotted on the histogram with bin size of 1,000. The mean and standard deviation of the bin counts are 1000.0±32.52, demonstrating the uniformity of biHash.

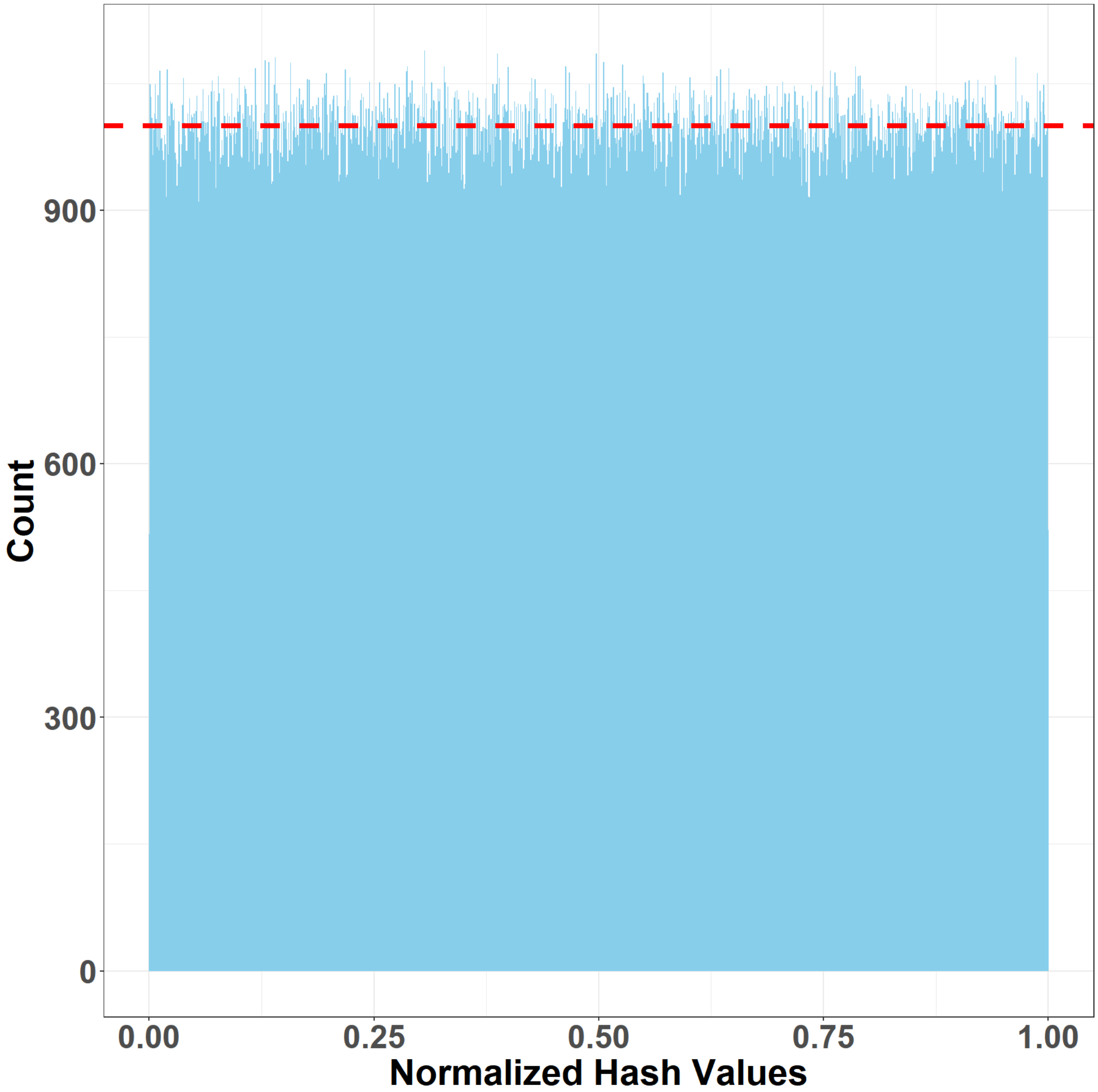

**Supplementary Figure S5. Histogram of 1,000,000 100-mer hashes generated by 2-base module of biHash from a random nucleotide sequence of length 1,000,099.** The red dashed line indicates the average number of hashes in a bin (1,000). The hash values were normalized by dividing them with $2^{64} - 1$, the largest 64-bit integer, and plotted on the histogram with bin size of 1,000. The mean and standard deviation of the bin counts are 1000.0±32.1, demonstrating the uniformity of biHash.

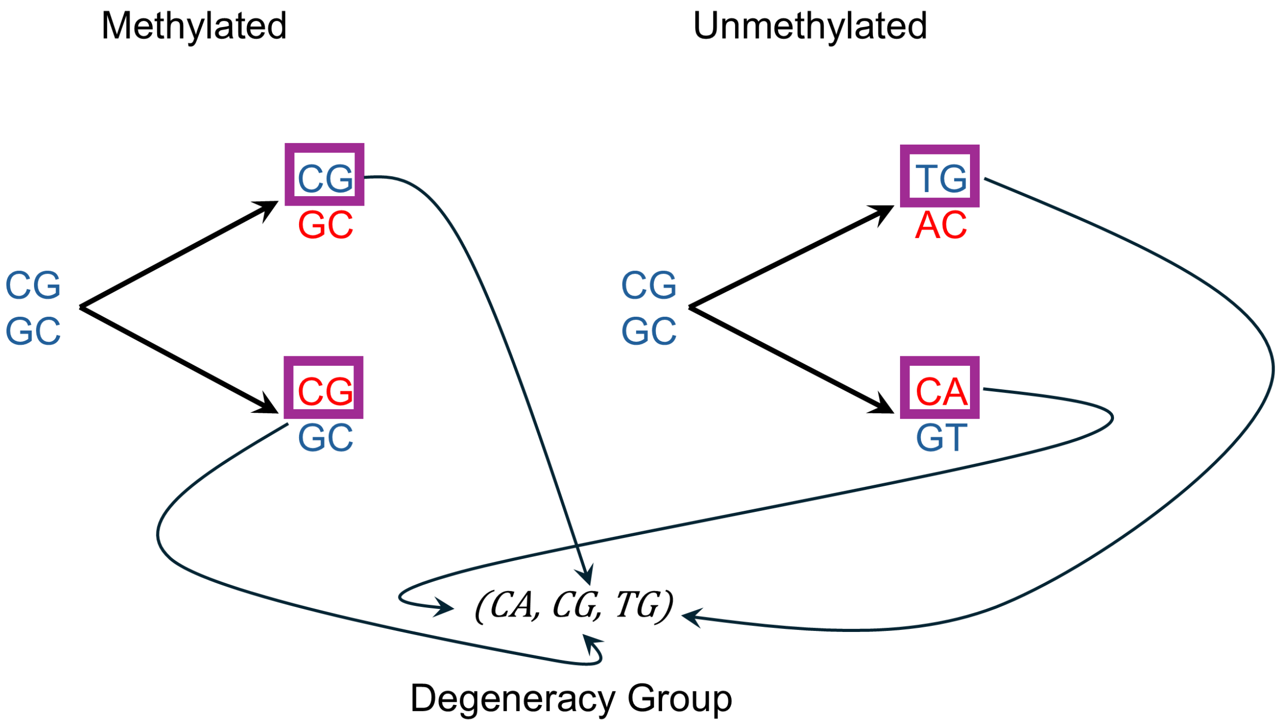

**Supplementary Figure S6. Symmetrical mapping of bisulfite-converted dinucleotides via dinucleotide alphabet degeneracy.** Original genomic DNA (blue) undergoes bisulfite treatment, where methylated cytosines remain intact and unmethylated cytosines are converted to uracil (read as thymine). During sequencing, both the forward strands and their respective reverse complements (red) are captured. Every resulting forward sequence variation (purple boxes) is mapped to a unified degeneracy group (CA,CG,TG).

**Supplementary Table S1. Hashing algorithms used to benchmark biHash.** All benchmarking tests were performed using 1 thread on a server-class system with 144 Intel(R) Xeon(R) Gold 6254 CPU @ 3.1 GHz with 2.9 TB RAM. Commit hashes are listed when there is no release version of the repository.

| Algorithm | Code repository | Version/commit |
| --- | --- | --- |
| City hash | https://github.com/google/cityhash | f5dc541 |
| Murmur hash | https://github.com/aappleby/smhasher | 92cf370 |
| xxHash | https://github.com/Cyan4973/xxHash | 0.8.1 |
| Rabin-Karp | https://github.com/TheAlgorithms/C-Plus-Plus | 23b133a |

**Supplementary Table S2. Kolmogorov–Smirnov (K-S) test statistics and associated *P*-values of biHash distributions.** Each module was tested against a theoretical uniform distribution using 1,000,000 normalized hash values from 100-mers of a simulated random nucleotide sequence.

| Modules | K-S statistic | *P*-value |
| --- | --- | --- |
| Dinucleotide-CG | 0.0008 | 0.62 |
| Dinucleotide-CA | 0.0008 | 0.62 |
| Dinucleotide-CC | 0.0007 | 0.65 |
| Dinucleotide-CT | 0.0005 | 0.95 |
| 3-base (C-to-T) | 0.0006 | 0.84 |
| 3-base (G-to-A) | 0.0006 | 0.83 |
| 2-base | 0.0006 | 0.89 |

**Supplementary Table S3. Shapiro–Wilk test P-values for the transformed $Z_0$ and $Z_1$ distributions of biHash modules.** As the normality test is sensitive to large sample sizes, we selected the first 1,000 transformed hash values, below SciPy's recommendation of 5,000, as input.

| Modules | $Z_0$ *P*-value | $Z_1$ *P*-value |
| --- | --- | --- |
| Dinucleotide-CG | 0.57 | 0.20 |
| Dinucleotide-CA | 0.16 | 0.87 |
| Dinucleotide-CC | 0.48 | 0.41 |
| Dinucleotide-CT | 0.79 | 0.30 |
| 3-base (C-to-T) | 0.10 | 0.73 |
| 3-base (G-to-A) | 0.36 | 0.67 |
| 2-base | 0.55 | 0.60 |

**Supplementary Table S4. Pearson correlation coefficients between the transformed $Z_0$ and $Z_1$ distributions for each biHash module.** Values are reported as r±SE, where SE represents one standard error.

| Modules | Pearson Correlation Coefficient |
| --- | --- |
| Dinucleotide-CG | $0.009 \pm 0.032$ |
| Dinucleotide-CA | $0.025 \pm 0.032$ |
| Dinucleotide-CC | $0.023 \pm 0.032$ |
| Dinucleotide-CT | $-0.019 \pm 0.032$ |
| 3-base (C-to-T) | $-0.011 \pm 0.032$ |
| 3-base (G-to-A) | $-0.010 \pm 0.032$ |
| 2-base | $0.031 \pm 0.032$ |

**Supplementary Table S5. Benchmarking script peak memory usage of each hashing algorithm generating 1 hash value for each *k*-mer with different *k*-mer lengths from 1,000,000 simulated sequences of length 250 nucleotides.**

| Algorithm | *k*-mer size | Peak memory usage (MB) |
|---|---|---|
| Dinucleotide-CG | 50 | 5.76 |
| Dinucleotide-CA | | 5.76 |
| Dinucleotide-CC | | 5.76 |
| Dinucleotide-CT | | 5.76 |
| 3-base (C-to-T) | | 5.76 |
| 3-base (G-to-A) | | 5.76 |
| 2-base | | 5.76 |
| CityHash | | 9.22 |
| xxHash | | 9.22 |
| MurmurHash | | 9.22 |
| Rabin-Karp | | 9.22 |
| Dinucleotide-CG | 150 | 5.76 |
| Dinucleotide-CA | | 5.76 |
| Dinucleotide-CC | | 5.76 |
| Dinucleotide-CT | | 5.76 |
| 3-base (C-to-T) | | 5.76 |
| 3-base (G-to-A) | | 5.76 |
| 2-base | | 5.76 |
| CityHash | | 9.22 |
| xxHash | | 9.22 |
| MurmurHash | | 9.22 |
| Rabin-Karp | | 9.22 |
| Dinucleotide-CG | 150 | 5.76 |
| Dinucleotide-CA | | 5.76 |
| Dinucleotide-CC | | 5.76 |
| Dinucleotide-CT | | 5.76 |
| 3-base (C-to-T) | | 5.76 |
| 3-base (G-to-A) | | 5.76 |
| 2-base | | 5.76 |
| CityHash | | 9.22 |
| xxHash | | 9.22 |
| MurmurHash | | 9.22 |
| Rabin-Karp | | 9.22 |
